## Supporting Text for "Horizontal transmission and recombination maintain forever young bacterial symbiont genomes"

#### Supporting Information

**Table of Contents:**

| | |
|---|---|
| P 1-5 | S1 Supporting Text: Supplemental Methods |
| P 5-16 | S2 Supporting Text: A Coalescent Model for Endosymbiont Populations with Mixed Transmission Modes |
| P 17-25 | S3 Supporting Text: Symbiont Species Descriptions |
| P 26-33 | Supporting Figures |
| P 34-41 | Supporting Tables |
| P 41-43 | Supporting References |

### S1 Supporting Text

#### Supplemental Methods

##### 1. Samples and Genomic Data Production, Continued

**Verification of species identification**

Species identifications for the bivalve samples were confirmed by BLAST. We extracted the full cytochrome oxidase 1 (CO1) gene sequence from each annotated mitochondrial genome to verify the host species. To verify the symbiont species/strain, we extracted the full 16S rRNA sequence from the annotated symbiont genomes. We then blasted both sequences against the non-redundant nucleic acid database from NCBI to find the closest sequence match.

We found that our *B. septemdierum* sequence is 99.87% similar to the other known *B. septemdierum* sequence (100% query coverage). Our *B. childressi* sequence too is nearly identical to the known *B. childressi* sequences, at 99.54-99.93% sequence identity (100% query coverage). Similarly, the symbionts of these two host species were most closely related at the 16S to the symbiont of *B. septemdierum* str. Myojin knoll (99.28% identity and 100% coverage) and *B. childressi* (99.53% identity and 98% coverage).

The *S. velum* samples were sequenced and identified previously [9], and the *S. pervernicosa* in that study came from the same sample set as the *S. pervernicosa* samples sequenced in this paper. Nevertheless, we compared the assembled symbiont 16S sequence against NCBI and found that the previously sequenced and assembled *S. pervernicosa* symbiont was the best hit (100% identity over 100% of query), and the other *S. pervernicosa* or *reidi* (synonym) sequences were >99.5% identical as well.

Although they were collected off of Oregon (JDF Ridge) and morphologically identified as *Calyptogena pacifica*, our *C. fausta* samples were identified as such by their CO1 sequence similarity of 97.32% to *C. fausta* in NCBI, as compared to a similarity of 91.63% to *C. pacifica* (both with 100% query coverage). Interestingly, the assembled symbiont 16S sequence was 100% identical to the *C. fausta* symbiont in NCBI (over 98% of the query) and it was also 99.86% identical to the *C. pacifica* symbiont (over 92% of the query). These two sequences were also very similar to each other (99.86% identical over 94% of the *C. fausta* sequence), indicating these symbionts are very closely related.

Lastly, we blasted the assembled *C. extenta* mitochondrial CO1 sequence to the NCBI database and found that this sample matches the NCBI sequences at 99.29-100% sequence identity (over amplicon fragments covering 33-42% of the query). Comparing the symbiont 16S sequence assembled from this sample to the database resulted in a hit to the C. extenta symbiont at 100% sequence identity (across 92% of the query).

##### Genotyping error rate estimation

Due to an inadvertent demultiplexing error for the single lane containing *C. fausta* samples, each sample was demultiplexed as a singly, i5, indexed sample. This led to an obvious increase in the rates of index hopping and therefore precluded confident analysis of within individual variation for these samples. Although we expect that consensus genome sequences would be largely unaffected, we decided to confirm this experimentally. We therefore prepared fresh libraries for four of these samples that we sequenced with other libraries and that were demultiplexed using both forward and reverse indexes (second coverage listed in S1 Table). In comparing the four symbiont genome assemblies with those from our initial effectively singly indexed run, we found only a single SNP difference across all four suggesting that our consensus assembly quality is in the range of Q67, or extremely accurate. We therefore used these four libraries for inter-individual variation analyses (S5 Table) and the original ten sequenced individuals for all consensus sequence-based analyses.

#### 2. Genome Analyses, Continued

##### Divergence dating

*Phylogeny inference and divergence date estimation, continued:*

To empirically assess what Beast2 divergence estimation parameters to use and how many calibration points to include, we first tested a range of settings run over a MCMC chain length of 100e6 using a HKY model of substitution and a random local molecular clock. We tested the results generated under Yule (birth-only) versus birth-death models of speciation with either a single calibration point at the root of Bivalvia (see Materials and Methods). We also tested multiple calibration points along the host tree for the nodes with fossil data available in the Fossilworks database [104] (i.e., Veneridae, Vesicomyidae, *Calyptogena,* Mytilidae, Ostreidae, Bathymodiolus, Modiolus, Mytilus, Crassostrea, and Solemya). To further assess the impact of node calibration on our results, we also tested uniform versus log-normal prior distributions for the node dates.

While an initial chain length of 100e6 was adequate for ascertaining that the single calibration point strategy was far superior to multiple calibration points in Beast2, the posterior distributions had not stabilized by this point (as visualized in Tracer [115]), so we ran the chains an order of magnitude longer, which produced well-sampled, converged prior distributions for all settings. We repeated these single calibration (Bivalvia or Gammaproteobacteria) runs for the four pairwise combinations of speciation model (Yule or birth-death) and prior node age distribution (uniform or lognormal) to ensure that the prior distributions converged on the same value when starting from a different seed. Trees were summarized as a maximum clade credibility tree with mean node heights in TreeAnnotator [103], with a 25% burn-in. While all parameter values resulted in similar results, the results from the Yule model with calibration dates modeled with log-normal distribution produced divergence dates most similar to expectations based on fossil evidence (see S12 Table).

We also inferred divergence times using maximum likelihood phylogenies and the RelTime relative rate framework implemented in MEGA [95,96]. This approach produced exceptionally similar values to Beast2, given that it took a fraction of the time to run and uncertainty in the tree was not considered. Given this speed, we were able to test multiple substitution models and models of rate heterogeneity (listed in S12 Table). RelTime's fast inference times also enabled us to estimate divergence dates for symbionts, which were problematic to infer in Beast2 (discussed in the Main Text). We used a single calibration point at the base of the ingroup, Gammaproteobacteria (see Materials and Methods). While some work has been done to infer bacterial divergence dates, few of the bacterial clades in our tree have sufficient evidence and/or genomic taxonomic sampling to warrant inclusion, so we did not test additional bacterial calibration points. In addition to RelTime, we also tried dating mitochondrial and symbiont maximum likelihood phylogenies in PATHd8 [69], which estimates node ages from the mean path lengths between nodes and leaves of these trees. However, PATHd8 did not perform nearly as well as RelTime for both hosts and mitochondria or Beast2 for mitochondria. See S12 Table for divergence date estimates from these different approaches and settings.

#### S2 Supporting Text

**A coalescent model for endosymbiont populations with mixed transmission modes**

##### Model Overview

We consider the coalescent process that relates a sample of **n** endosymbiont lineages. Briefly, endosymbiont populations are structured such that all are contained within sub-populations, an "infection" within a single host individual. It is assumed that all symbiont reproduction occurs within a host and that there is a constant rate of horizontal transfer, **H**, between hosts every host generation. In this way, the hosts, numbering $\mathbf{N_H}$ are similar to demes within classical metapopulation models (*e.g.*[116–118]). Hosts are related to one another following a Wright-Fisher model with constant population size and are exchangeable within the model. For simplicity, with the exception of within-host processes, we measure time in host generations. There are therefore three possible states for a pair of symbiont lineages. They may be (1) occupying different host infections, (2) occupying the same host infection and (3) coalesced (S5 Fig.).

**Transition Rates and Host Coalescence**

Here, we consider the coalescent process for a sample of two symbiont lineages. Under this model, the probability of a horizontal transmission event that brings a symbiont lineage from one host maternal lineage into another that contains the other symbiont lineage is then

$$p(hori\_trans) = \frac{2H}{N_H} \qquad \text{Equation 1}$$

Here we assume that the H is sufficiently small that H^2 terms can be safely ignored. If we don't make this assumption, this probability would instead be (2H-H^2)/NH. We note that due to the exchangeability of the host lineages, we can ignore all horizontal transmission events that do not bring two symbiont lineages into a single host. In the context of this model, those events are unobservable. However, two lineages may also transition from two distinct hosts to the same in the previous host generation due to coalescence among the host maternal lineages as well.

$$p(host\_coal) = \frac{1}{N_H} \qquad \text{Equation 2}$$

giving a total probability of two lineages transitioning from two separate host infections into a single host during a single host generation.

$$p(same\_host) = \frac{2H+1}{N_H}$$ Equation 3

This model excludes the possible contribution of male host lineages to coalescence within symbiont populations. If the horizontal transmission probability is very high, this assumption might not be realistic because symbiont lineages could trace their ancestry through males so long as they transmit horizontally within a single host generation. However, if the horizontal transmission rate is relatively low this can be ignored because males will not transmit symbiont populations to their progeny and the majority of sampled symbiont lineages will trace their ancestry almost entirely through maternal host lineages..

**Within Host Symbiont Dynamics**

We consider that each host begins with $\mathbf{N_S}$ symbiont individuals provisioned by the mother. *I.e.*, this is the size of the transmission bottleneck between host generations. The infection then grows following binary growth for a number of generations $\mathbf{G_S}$. Therefore there are $\mathbf{G_S}$ symbiont generations per host generation. The probability of coalescence per symbiont generation i in $\{1 \ldots G_S\}$

$$p(coal_i) = \frac{1}{2^{i-1} N_S}$$ Equation 4

for two symbiont lineages within the same host individual that are not coalesced. This ignores the possibility of coalescence after population growth has finished. However, symbiont population sizes within hosts are typically on the order of billions [119–124], and the probability of coalescence is negligible after the first few generations of doubling growth. Therefore, the total probability of symbiont coalescence per host generation for two symbiont lineages that are within the same host, can be approximated as

$$p(coal) = 1 - \prod_{i=1}^{G_s} 1 - \frac{1}{2^{i-1} N_s} \quad \text{Equation 5}$$

We note that for larger values of Ns, this can be approximated as 2/Ns quite well. In the absence of a horizontal transmission event or a coalescent event, both symbiont lineages are transmitted to the host's mother in the previous generation. There is no specific requirement for binary growth within the symbiont population and this process could be modeled more generally using an arbitrary vector of symbiont population sizes if information about symbiont population size dynamics is known (*e.g.* by sampling tissues during development as in [31,33]). However, the smallest symbiont population size will tend to dominate patterns of within host coalescence and therefore estimating the transmission bottleneck inoculate size is likely to be the most pertinent for generating realistic models of symbiont coalescence. For our purposes here, the rate is not particularly consequential.

Alternatively, given that two symbiont lineages are in a single host but have not coalesced, the probability of that one transitions out of the host infection into another host infection is

$$p\left(diff\_host\right) = \frac{2H(N_H\text{-}1)}{N_H}$$ Equation 6

Therefore, given that two lineages are in the same host, the probability that they coalesce prior to transmitting out of that host maternal lineage is

Equation 7
$$p\left(coal \mid same\_host\right) = \frac{p\left(coal\right)}{p\left(coal\right) + p\left(transmit\_out\right)}$$

**Time to Coalescence**

In this model, and consistent with most symbiont datasets we are aware of, we consider the case where two symbiont lineages are sampled from different hosts. The number of times that one of the two lineages will enter the same host as the other is geometrically distributed with mean 1/p(coal|same_host), and the expected time until they coalesce, measured in host generations, is the product of the expected number of these events that occur prior to coalescence multiplied by the expected time for each event that brings the two lineages into the same host and they either coalescences or one lineage moves horizontally into another host infection, *i.e.*,

$$E[t_{coal}] = \frac{1}{p\left(coal \mid same\_host\right)}\left(\frac{1}{p\left(same\_host\right)} + \frac{1}{p\left(coal\right) + p\left(transmit\_out\right)}\right)$$ Equation 8

Assuming a Poisson distributed per symbiont generation mutation rate μ and an infinite sites model, we obtain the expected number of mutations that separate the two symbiont lineages as

$$E[S] = E[t_{coal}]2\mu G_S$$ Equation 9

Below, we demonstrate that under some conditions these populations can be modeled using Kingman's coalescent, but the above results should hold regardless. Although it is not the primary goal of this work, we also confirmed these predictions using forward-in-time simulations over a range of feasible population conditions (S6 Fig.). This indicates that these equations can correctly predict the nucleotide diversity and time to coalescence for two lineages under arbitrary models of symbiont evolution including those where $N_H$ is of the same order as $N_S$. Our software to simulate this in forward time is available from https://github.com/shelbirussell/ForeverYoungGenomes_Russell-et-al, but we note that it will generally be much more efficient to use coalescent simulation so long as the population can be modelled appropriately within that framework.

**Kingman's coalescent**

If the host population size and symbiont inoculate size, $N_H$ and $N_S$, are of similar magnitude, this model cannot be approximated using Kingman's coalescent and $t_{coal}$ will not be exponentially distributed but rather it will be a mixture of rates. However, for many endosymbiont populations of interest, $N_H$ is likely to be much larger than $N_S$ [120,125–128]. Therefore, the primary rate limiting process for coalescence will be the time until two symbiont lineages are found within the same host maternal lineage, a result closely related to those from metapopulation theory (*e.g.*, [116–118]). Using the framework laid out above, the exact, single host-generation transition matrix, **P,** with probability of coalescence given two lineages are within the same host in the same host generation, **C**, for states (1) two symbiont lineages in different host infections, (2) two symbiont lineages in the same host infection, and (3) coalesced, can be expressed as

$$P = \begin{bmatrix} 1-\frac{(2H+1)}{N_H} & \frac{(2H+1)}{N_H} & 0 \\ \frac{2H(N_H-1)}{N_H} & 1-\frac{2H(N_H-1)}{N_H}-C & C \\ 0 & 0 & 1 \end{bmatrix}$$

Equation 10

We now use the result of Möhle [129,130] to evaluate convergence in Markov processes that operate on two distinct time scales. The above transition matrix, P, contains terms that become increasingly different in the limit as $N_H$ approaches infinity.  This matrix can be subdivided into two classes of events. Specifically, F, contains the so-called “fast” events that do not depend on $N_H$. S contains the “slow” events that depend on the reciprocal of $N_H$. Finally, an additional term, o(1/N), may contain the terms that trend to zero more quickly than $1/N_H$, however here, this term is equal to zero and omitted below.

$$F = \begin{bmatrix} 1 & 0 & 0 \\ 2\mathrm{H} & 1-2H\text{-C} & C \\ 0 & 0 & 1 \end{bmatrix}$$

$$S = \begin{bmatrix} \text{-2H-1} & (2\mathrm{H}+1) & 0 \\ 0 & 0 & 0 \\ 0 & 0 & 0 \end{bmatrix}$$ Equation 11

Then, if the matrix E = limit(t->infinity)F^t exists, in the continuous approximation of the ancestral process, all of the fast events reach their equilibrium immediately. This can be expressed as

$$E = \begin{bmatrix} 1 & 0 & 0 \\ \frac{2H}{2H+C} & 0 & \frac{C}{2H+C} \\ 0 & 0 & 1 \end{bmatrix}$$ Equation 12

This can be interpreted as follows. Consider a case where two lineages are sampled from the same host (row 2), they will either coalesce or horizontally transfer into two separate host infections essentially immediately with probability equivalent to the relative rates of those possible outcomes. Conversely, if two lineages are already in two separate hosts, they will remain there indefinitely on the time-scale of the “fast” events.

Then, by taking the matrix product, ESE, we obtain the rate matrix G, which in combination with E describes the rescaled ancestral process of the two samples, with time now measured in units of $N_H$ generations. This follows from Möhle's [129,130] theorem, which formally states that this rescaled ancestral process converges to $\lim_{n\to\infty} P^{Nt} = Eexp(tG)$. Below is the rate matrix, G

$$G = \begin{bmatrix} -1-2H+\frac{2H(1+2H)}{C+2H} & 0 & \frac{C(1+2H)}{C+2H} \\ \frac{4H^2(1+2H)}{(C+2H)^2}+\frac{2(-1-2H)H}{C+2H} & 0 & \frac{2CH(1+2H)}{(C+2H)^2} \\ 0 & 0 & 0 \end{bmatrix}$$

Equation 13

This result implies that when two lineages are sampled from different host infections, the rate of coalescence could be further rescaled by (C+2H)/(C(1+2H)), and for most of the plausible parameter space, the total symbiont effective population size will exceed $N_H$. For example, if C = 0.02, H=0.01, and $N_H$ = 10,000, the symbiont effective population size would be 19,608. This indicates that effective population sizes may sometimes be substantially larger than $N_H$ and is consistent with metapopulation coalescent theory with moderate barriers to migration among small demes [116,117].

**Extension to Larger Samples**

For the Kingman coalescent to be a valid approximation for evolution of endosymbiont lineages, coalescent events among lineages must be primarily pairwise rather than multiple mergers among several lineages [131–133]. Given the complexity of this model, it is not feasible to evaluate a three or four lineage exact backwards in time transition matrix as we did for two lineages above.

Instead, we make use of the following heuristic observation. In order for symbiont lineages to experience a multiple coalescent event there must be three or more symbiont lineages within a single host at one time. Because we assume that all symbiont lineages are sampled from distinct hosts. Conditional on two lineages occupying the same infection, they will either coalesce or one migrate out with probability p(coal or leave) = C+2H. Conversely, another lineage will enter the same infection with probability p(enter) = (1+H)/$N_H$. In the limit as $N_H$ trends towards infinity, this value rapidly approaches zero. Because coalescence or departure from the same infection occurs on a much faster time-scale than migration events, all coalescent events must be pairwise in the limiting case, where higher order coalescent events would occur at a rate on the order of 1/$N_H$^2.

**Modeling Recombination**

It is increasingly widely appreciated that recombination can have a profound influence on bacterial populations [44,45,134,135]. Throughout, we model recombination as equivalent to gene conversion in eukaryotic species (following *e.g.,* [111]). When a gene conversion event occurs on a genealogy, it creates an additional lineage within the converted segment to the right and to the left of the recombination breakpoints. Initially, these two symbiont lineages will be present within the same infection and if they coalesce before either lineage transmits into another host maternal lineage, this event will not be observed. Therefore, the observable recombination rate is decreased relative to the per generation recombination rate. Thus, the probability that the recombination event is unobserved due to coalescence within the same infection is p(unobserved) = C/(C+2H).

There is also a chance that two lineages coalesce prior to coalescing with any other lineages due to subsequent back-migration into the same host maternal lineage, and subsequent coalescence. This would also cause a recombination event to be unobserved and is already an integral part of most coalescent simulation algorithms (*e.g.*, [136,137]), which model the probability of back coalescence to the marginal genealogy to the left. For our purposes it is therefore sufficient to multiply the per-generation recombination rate by a constant factor of 2H/(C+2H) to obtain a corrected effective recombination rate estimate in the Kingman coalescent. Importantly, when horizontal transmission is rare relative to within-host coalescence, the effective recombination rate will tend to be quite small for symbiont populations.

**Possible Extensions Using Scattering Phase Models**

One key implication of this model is that when multiple symbiont lineages are sampled from within a single host, they may sometimes coalesce quickly before either symbiont lineage horizontally transfers into another host maternal lineage. This could feasibly be extended using a separation of time-scales approach as in the “scattering” and “collecting” phases of metapopulation coalescent theory [116–118]. However, we do not believe this addition is necessary for the interpretation of most existing endosymbiont datasets. In general, two lineages that coalesce very quickly will have few polymorphisms that distinguish them. Given current sequencing practices it may be challenging to infer the presence of multiple lineages unless one is a very recent migrant—obviating this concern—and suggesting that these considerations will not significantly affect interpretations of empirical endosymbiont genomic diversity datasets. Nonetheless, as microbial sampling practices continue to progress, with *e.g.* single-cell sequencing approaches becoming more common, modeling these additional considerations may be important for future applications of this framework that do sample multiple lineages from within the same host. However in the analyses presented here, unless otherwise stated, we will sample on a single symbiont lineage from each host individual.

**Summary and Assumptions**

We have derived a coalescent model for endosymbiont lineages that are partially structured by host populations and experience both horizontal and vertical transmission. We stress that, as with any theoretical framework, there are many assumptions underlying this result and that these assumptions should be critically evaluated when applying these to additional populations and datasets. In particular, this model explicitly assumes that the host population is at a demographic equilibrium and neutrally evolving. Furthermore, we assume that the rate of horizontal transmission is relatively small and that the host population size is much larger than the within-host symbiont effective population size. Although extensions are possible to accommodate these and other added complexities, they are beyond the scope of our treatment here where we focus primarily on understanding the evolution of the endosymbiont populations during mixed transmission modes. However, importantly for this work, this model justifies our use of standard Kingman coalescent simulation software, as it should approximate the evolution of endosymbiont populations during mixed transmission modes and provides a theoretical justification for the mechanistic relationships between horizontal transmission and observable recombination events that have the potential to create genetically diverse symbiont chromosomes.

#### S3 Supporting Text

##### Symbiont Species Descriptions

**Name:** *Candidatus* Thiodubiliella gen. nov.

Thī.ō.dŭb.ĭlĭ.ĕllă N.L. adj. *thio* sulfur*;* N.L. n. *dubiliella* named for Prof. Nicole Dubilier who has worked on the bathymodiolin deep-sea symbioses throughout her career and contributed significant knowledge to the field about their biology; N.L. fem. n. *Thiodubiliella.*

**Properties:** Bacterial symbionts in the SUP05 clade of Gammaproteobacteria that associate with deep-sea mytilid mussels at hydrothermal vents. Short, rod-shaped bacteria (~0.6 µm long) live intracellularly within specialized host gill cells [137]. Their genomes encode genes for sulfide and hydrogen oxidation (see *Bathymodiolus septemdierum* symbiont from Lau genome annotation under NCBI BioProject number PRJNA562081 and [42]). Genus diagnosed molecularly by phylogenetic divergence (Figure 5A) and high sequence identity (>95%) at rRNA, core metabolic, and housekeeping genes. Ideally, average nucleotide identity across the genome is >73% for members of the genus (as in [138]). However, if high sequence identity at conserved elements is observed, this constraint can be relaxed (because symbiotic genomes often have increased structural dynamics due to bouts of inefficient selection). Genome structure is indeed dynamic in this clade, leading to much higher divergence at the genome level than the corresponding 16S rRNA level (*e.g.*, 33.1% and 98.6% identity, respectively, between *Ca.* Thiodubiliella symbionts from *Bathymodiolus septemdierum* (Lau) and the unnamed symbionts from *Bathymodiolus thermophilus* (NCBI accession GCF_003711265.1)). The genus exhibits approximately 35% and 95% sequence similarity at the genome an 16S levels, respectively, to *Candidatus* Thioglobus autotrophicus (unclassified Gammaproteobacteria; NCBI accession GCF_001293165.1) in the closest related genus (see Figure 5A).

**Typification:** Class: Gammaproteobacteria; Order: unclassified.

**Type species:** *Candidatus* Thiodubiliella endoseptemdiera sp. nov.

**Name:** *Candidatus* Thiodubiliella endoseptemdiera sp. nov.

ĕn.dō.sĕpt.ĕm.dĭēr.ă Gr. fem. adj. *endo* within; N.L. n. *septemdiera* (host) *Bathymodiolus septemdierum*; N.L. fem. n. *endoseptemdiera*.

**Properties:** Intracellular thioautotrophic bacterial gill symbiont of bathymodiolin mussels. Species affiliation defined molecularly by divergence (Figure 5), by ≥95% average nucleotide identity genome-wide, and ≥98.7% sequence identity at rRNA, core metabolic, and housekeeping genes in the genome (*Bathymodiolus septemdierum* symbiont genome annotation under NCBI BioProject number PRJNA562081).

**Type host:** *Bathymodiolus septemdierum* (Bivalvia: Pteriomorphia: Mytilidae).

**Other hosts:** None.

**Type locality:** Tu'i Malila, Lau Basin, S21° 59.3455′ W176° 34.0926′.

**Other localities:** None.

**Specimens deposited:** None, as this bacterium is currently unculturable and is only known molecularly and microscopically. This is a tentative name proposed with the designation "*Candidatus*".

**Name:** *Candidatus* Methanofisheria gen. nov.

Mĕth.ănō.fĭsh.ĕr.ă N.L. adj. *Methano* methane; N.L. n. *fishera* named for Prof. Charles Fisher who has worked on the bathymodiolin deep-sea symbioses throughout his career and contributed significant knowledge to the field about their biology; N.L. n. fem. *Methanofisheria*.

**Properties:** Bacterial symbionts in a clade of methane-oxidizing Gammaproteobacteria that associate with deep-sea mytilid mussels at hydrothermal vents and cold seeps.

Coccoid-shaped bacteria (1.4-2 μm in diameter) containing stacked membranes live intracellularly within specialized host gill cells [139]. Their genomes encode pathways for using methane as both a carbon and an energy source (see *Bathymodiolus childressi* symbiont genome annotation under NCBI BioProject number PRJNA562081), and hosts have been shown to uptake methane [140]. Genus diagnosed molecularly by phylogenetic divergence (Figure 5A) and high sequence identity (>95%) at rRNA, core metabolic, and housekeeping genes. Ideally, average nucleotide identity across the genome is >73% for members of the genus (as in [138]). However, if high sequence identity at conserved elements is observed, this constraint can be relaxed (because symbiotic genomes often have increased structural dynamics due to bouts of inefficient selection). Genome structure is indeed dynamic in this clade, leading to much higher divergence at the genome level than the corresponding 16S rRNA level (*e.g.*, 58.3% and 97.8% identity, respectively, between *Ca.* Methanofisheria symbionts from *Bathymodiolus childressi* and the unnamed symbionts from *Bathymodiolus platifrons* (NCBI accession GCF_002189065.1)). The genus exhibits approximately 50.2% and 94.0% sequence similarity at the genome and 16S levels, respectively, to *Candidatus* Methylobacter oryzae (Gammaproteobacteria: Methylococcales: Methylococcaceae; NCBI accession GCF_003994235.2) in the closest related genus (see Figure 5A).

**Typification:** Class: Gammaproteobacteria; Order: unclassified.

**Type species:** *Candidatus* Methanofishera endochildressiae sp. nov.

**Name:** *Candidatus* Methanofishera endochildressiae sp. nov.

ĕn.dō.chíld.rĕs.sĭ.āĕ Gr. adj. *endo* within; N.L. n. *childressiae* (host) *Bathymodiolus childressi*.

**Properties:** Intracellular methanotrophic bacterial gill symbiont of bathymodiolin mussels. Species affiliation defined molecularly by divergence (Figure 5), by ≥95% average nucleotide identity genome-wide, and ≥98.7% sequence identity at rRNA, core metabolic, and housekeeping genes in the genome (*Bathymodiolus childressi* symbiont genome annotation under NCBI BioProject number PRJNA562081).

**Type host:** *Bathymodiolus childressi* (Bivalvia: Pteriomorphia: Mytilidae)

**Other hosts:** None.

**Type locality:** Veatch Canyon, 39.8061°N 69.5922°W

**Other localities:** None.

**Specimens deposited:** None, as this bacterium is currently unculturable and is only known molecularly and microscopically. This is a tentative name proposed with the designation "*Candidatus*".

**Name:** *Candidatus* Cavanaughia gen. nov.

Cāv.ān.ăŭgh.ĭ.ă N.L. n. fem. named for Prof. Colleen Cavanaugh, who was one of the first to discover chemosynthetic symbiosis and worked on the *S. velum* symbiosis throughout her career and contributed significant knowledge to the field about their biology.

**Properties:** Gammaproteobacterial symbionts that associate with burrowing solemyid bivalves that inhabit continental shelf sediments. Rod-shaped bacteria (2-10 µm long) live intracellularly within specialized host gill cells [141]. Their genomes encode genes for sulfide oxidation (see [13]), and these associations have been shown to oxidize sulfide and fix carbon dioxide [141]. Genus diagnosed molecularly by phylogenetic divergence (Figure 5A) and high sequence identity (>95%) at rRNA, core metabolic, and housekeeping genes. Ideally, average nucleotide identity across the genome is >73% for members of the genus (as in [138]). However, if high sequence identity at conserved elements is observed, this constraint can be relaxed (because symbiotic genomes often have increased structural dynamics due to bouts of inefficient selection). Genome structure is indeed dynamic in this clade, leading to much higher divergence at the genome level than the corresponding 16S rRNA level (*e.g.*, 51.6% and 99.7% identity, respectively, between *Ca.* Cavanaughia symbionts from *Solemya velum* and *Solemya elarraichensis* (NCBI accession GCF_002021095.1)). The genus exhibits approximately 39.9% and 93.1% sequence similarity at the genome an 16S levels, respectively, to *Thiolapillus brandeum* (unclassified Gammaproteobacteria; NCBI accession GCF_000828615.1) in the closest related genus (see Figure 5A).

**Typification:** Class: Gammaproteobacteria; Order: unclassified.

**Type species:** *Candidatus* Cavanaughia endovela sp. nov.

**Name:** *Candidatus* Cavanaughia endovela sp. nov.

ĕn.dō.vē.lă Gr. adj. *endo* within; N.L. *vela* (host) *Solemya velum;* N.L. n. fem. *endovela*.

**Properties:** Intracellular thioautotrophic bacterial gill symbiont of *Solemya velum* bivalves. Species affiliation defined molecularly by divergence (Figure 5), by ≥95% average nucleotide identity genome-wide, and ≥98.7% sequence identity at rRNA, core metabolic, and housekeeping genes in the genome (NCBI accession GCF_000787395.1; [13]).

**Type host:** *Solemya velum* (Bivalvia: Protobranchia: Solemyidae)

**Other hosts:** None.

**Type locality:** Naushon Island, Woods Hole, MA, USA

**Other localities:** intertidal-subtidal sediments along the eastern North American coast off: Kingston Bay, Duxbury, MA (42°0'14.15"N, 70°40'43.79"W); Point Judith, Narragansett, RI

(41°22'55.91"N, 41°22'55.91"N); Shark River Island, Neptune Township, NJ (40°11'9.60"N, 74° 1'48.00"W); Sinepuxent Bay, MD (38°14'58.56"N, 75°9'8.06"W); Pivers Island, Beaufort, NC (34°42'59.23"N, 76°40'26.33"W).

**Specimens deposited:** None, as this bacterium is currently unculturable and is only known molecularly and microscopically. This is a tentative name proposed with the designation "*Candidatus*".

**Name:** *Candidatus* Reidiella gen. nov.

Rēĭd.ĭ.ĕl.lă N.L. n. fem. named for Prof. Robert Reid who first reported on these mouthless and gutless protobranch bivalves (from the synonymized *S. reidi*).

**Properties:** Gammaproteobacterial symbionts that associate burrowing solemyid bivalves that inhabit continental shelf sediments. Rod-shaped bacteria (2-5 μm long) live intracellularly within specialized host gill cells [30]. Their genomes encode genes for sulfide oxidation (see *Solemya pervernicosa* symbiont genome annotation under NCBI BioProject number PRJNA562081), and these associations have been shown to oxidize sulfide and fix carbon dioxide [142]. Genus diagnosed molecularly by phylogenetic divergence (Figure 5A) and high sequence identity (>95%) at rRNA, core metabolic, and housekeeping genes. Ideally, average nucleotide identity across the genome is >73% for members of the genus (as in [138]). However, if high sequence identity at conserved elements is observed, this constraint can be relaxed (because symbiotic genomes often have increased structural dynamics due to bouts of inefficient selection). Genome structure is indeed dynamic in this clade (Figure 4), however, close relatives of the *Ca.* Reidella symbiont from *S. pervernicosa* have not been sampled. Compared to one of the most closely related species, *Candidatus* Tenderia electrophaga (NCBI accession

GCA_001447805.1), *Ca.* Reidella endopervernicosa is 30.4% and 93.50% identical at the whole genome and 16S levels, respectively.

**Typification:** Class: Gammaproteobacteria; Order: unclassified.

**Type species:** *Candidatus* Reidiella endopervernicosa sp. nov.

**Name:** *Candidatus* Reidiella endopervernicosa sp. nov.

ĕn.dō.per.ver.ni.cō.sa Gr. adj. *endo* within; N.L. n. *pervernicosa* (host) *Solemya pervernicosa*; N.L. n. fem. *endopervernicosa.*

**Properties:** Intracellular thioautotrophic bacterial gill symbiont of *Solemya pervernicosa* bivalves. Species affiliation defined molecularly by divergence (Figure 5), by ≥95% average nucleotide identity genome-wide, and ≥98.7% sequence identity at rRNA, core metabolic, and housekeeping genes in the genome (*Solemya pervernicosa* symbiont genome annotation under NCBI BioProject number PRJNA562081).

**Type host:** *Solemya pervernicosa* (Bivalvia: Protobranchia: Solemyidae)

**Other hosts:** syn. *Solemya reidi.*

**Type locality:** sewage outfall, off Santa Monica, CA, USA

**Other localities** off Cape Nomamisaki, Kagoshima, Japan (NCBI accession AB499617).

**Specimens deposited:** None, as this bacterium is currently unculturable and is only known molecularly and microscopically. This is a tentative name proposed with the designation "*Candidatus*".

**Name:** *Candidatus* Ruthia endofausta sp. nov.

ĕn.dō.făŭ.stă Gr. adj. *endo* within; N.L. n. *fausta* (host) *Calyptogena fausta;* N.L. n. fem. *endofausta*.

**Properties:** Intracellular thioautotrophic bacterial gill symbiont of *Calyptogena fausta* clams. Species affiliation defined molecularly by divergence (Figure 5 and [24]), by ≥95% average nucleotide identity genome-wide, and ≥98.7% sequence identity at rRNA, core metabolic, and housekeeping genes in the genome (*Calyptogena fausta* symbiont genome annotation under NCBI BioProject number PRJNA562081). These criteria place the *Ca.* Ruthia endofausta symbiont in the "group 2" vesicomyid symbiont clade from [24]. Furthermore, these criteria distinguish this species from the type species for the "group 2" genus *Candidatus* Ruthia, *Ca.* Ruthia magnifica (NCBI accession GCF_000015105.1)*,* which is 66.8% identical genome wide and 97.9% identical at the 16S rRNA to *Ca.* Ruthia endofausta. As a member of the SUP-05 clade of sulfur-oxidizing bacteria, *Ca.* Ruthia endofausta is 42.7% and 95.8% identical at genome and 16S rRNA levels, respectively, to *Ca.* Thioglobus autotrophicus (NCBI accession GCF_001293165.1) in the closest-related genus.

**Type host:** *Calyptogena fausta* (Bivalvia: Heterodonta: Vesicomyidae)

**Other hosts:** None.

**Type locality:** Juan De Fuca Ridge, OR, USA.

**Other localities:** None.

**Specimens deposited:** None, as this bacterium is currently unculturable and is only known molecularly and microscopically. This is a tentative name proposed with the designation "*Candidatus*".

**Name:** *Candidatus* Vesicomyosocius endoextente sp. nov.

ĕn.dō.ĕx.tĕn.tĕ Gr. adj. *endo* within; N.L. n. *extenta* (host) *Calyptogena extenta*; N.L. n. masc. *endoextente*.

**Properties:** Intracellular thioautotrophic bacterial gill symbiont of *Calyptogena extenta* clams. Species affiliation defined molecularly by divergence (Figure 5 and [24]), by ≥95% average nucleotide identity genome-wide, and ≥98.7% sequence identity at rRNA, core metabolic, and housekeeping genes in the genome (*Calyptogena extenta* symbiont genome annotation under NCBI BioProject number PRJNA562081). These criteria place the *Ca.* Vesicomyosocius endoextente symbiont in the "group 1" vesicomyid symbiont clade from [24]. Furthermore, these criteria distinguish this species from the type species for the "group 1" genus *Candidatus* Vesicomyosocius, *Ca.* Vesicomyosocius okutanii (NCBI accession GCF_000010405.1), which is 88.2% identical genome wide and 99.2% identical at the 16S rRNA to *Ca.* Vesicomyosocius endoextente. As a member of the SUP-05 clade of sulfur-oxidizing bacteria, *Ca.* Vesicomyosocius endoextente is 37.9% and 94.4% identical at genome and 16S rRNA levels, respectively, to *Ca.* Thioglobus autotrophicus (NCBI accession GCF_001293165.1) in the closest-related genus.

**Type host:** *Calyptogena extenta* (Bivalvia: Heterodonta: Vesicomyidae)

**Other hosts:** None.

**Type locality:** Monterey Canyon, CA 36-40.92N 122-7.21W

**Other localities:** None.

**Specimens deposited:** None, as this bacterium is currently unculturable and is only known molecularly and microscopically. This is a tentative name proposed with the designation "*Candidatus*".

#### Supporting Figures

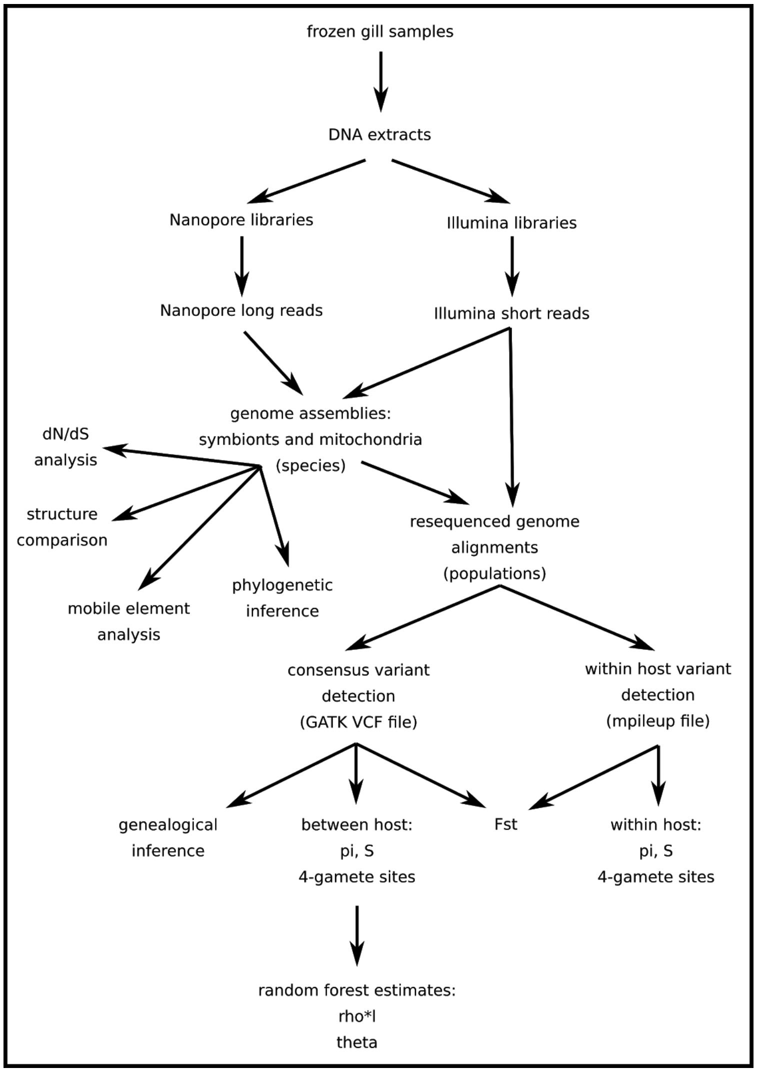


**S1 Fig.** Overview of the genomic data production and analysis steps used to study the population genomic processes influencing endosymbiont genome evolution.

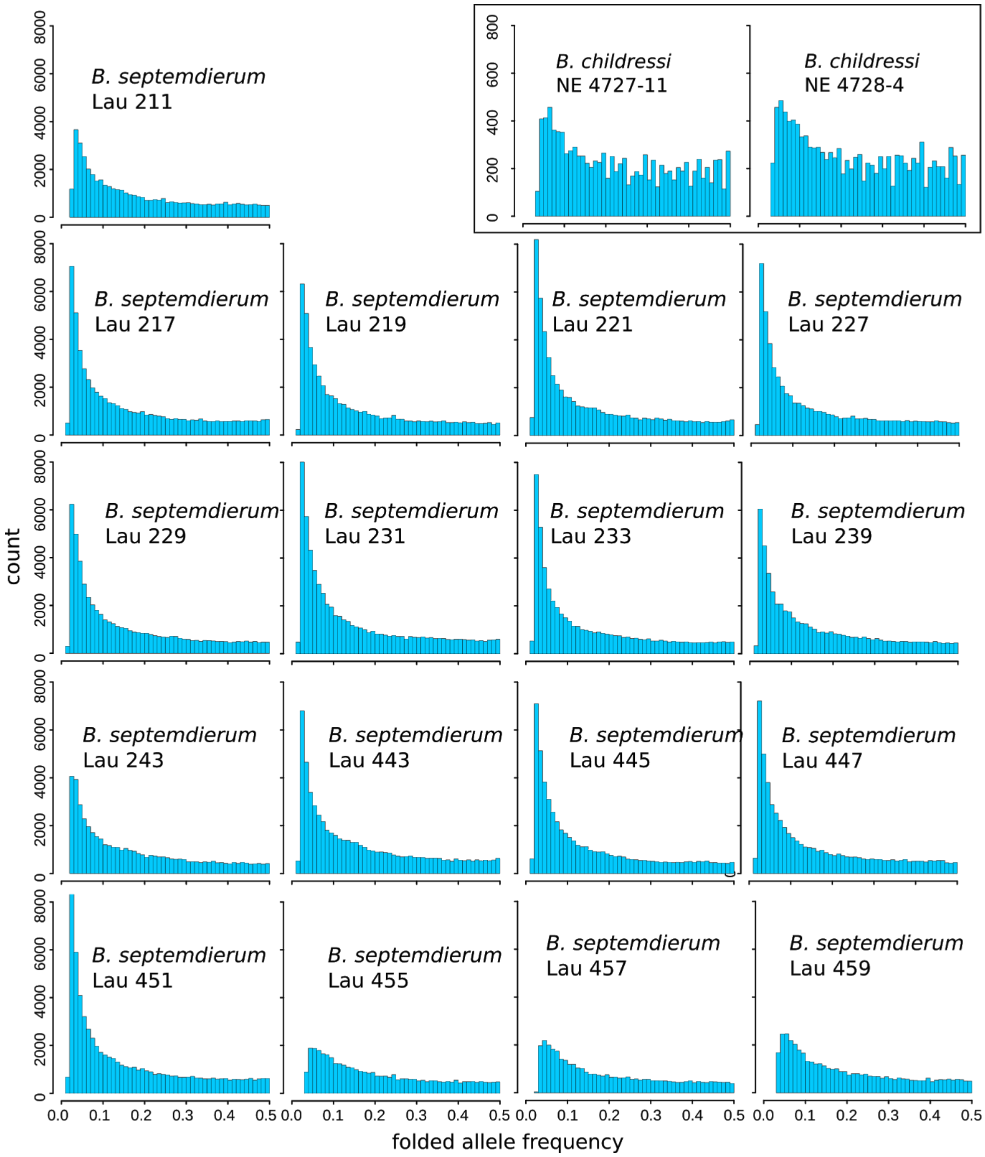


**S2 Fig.** Within-host symbiont folded allele frequency spectra for all *B. septemdierum* and *B. childressi* intrahost samples with more than 50x and 45x Illumina sequencing coverage, respectively (see S1 Table for coverages and S5 Table for diversity statistics).

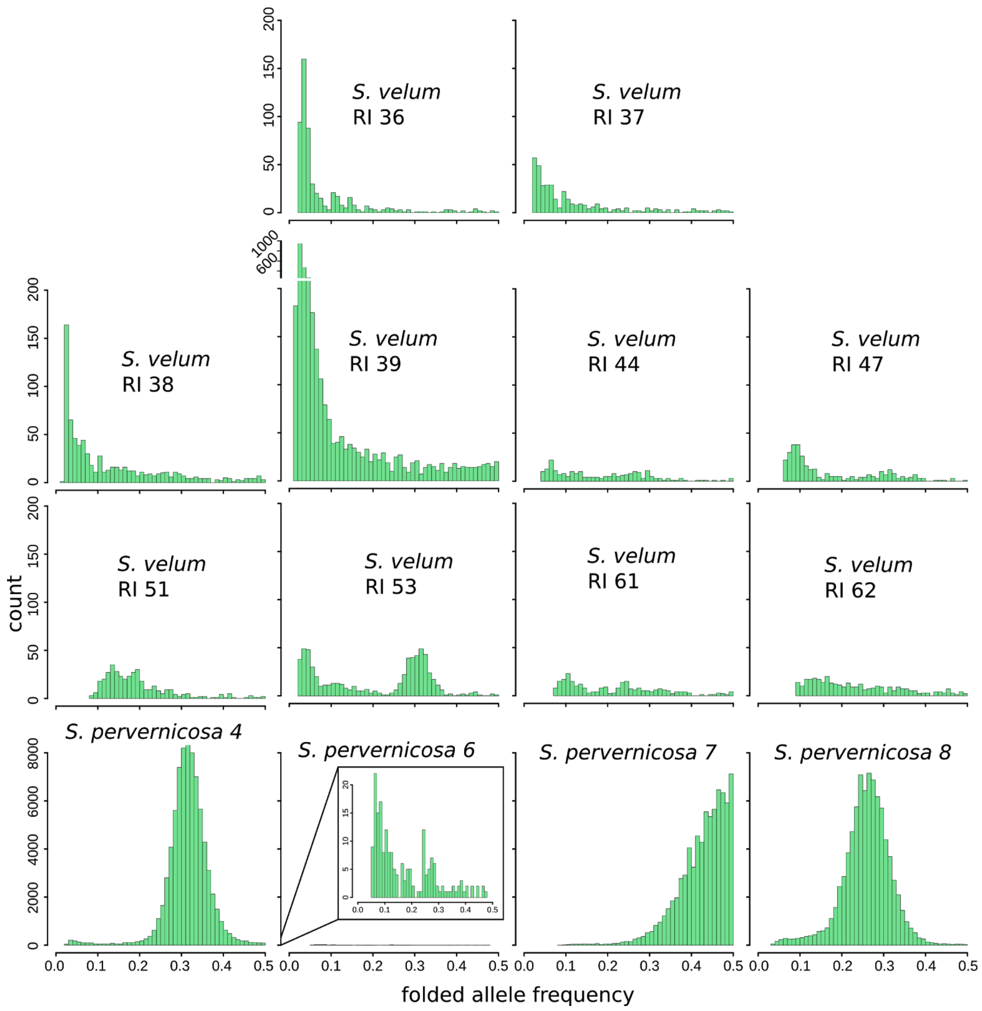


**S3 Fig.** Within-host symbiont folded allele frequency spectra for all *Solemya velum* and *Solemya pervernicosa* intrahost samples with more than 50x Illumina sequencing coverage (see S1 Table for coverages and S5 Table for diversity statistics).

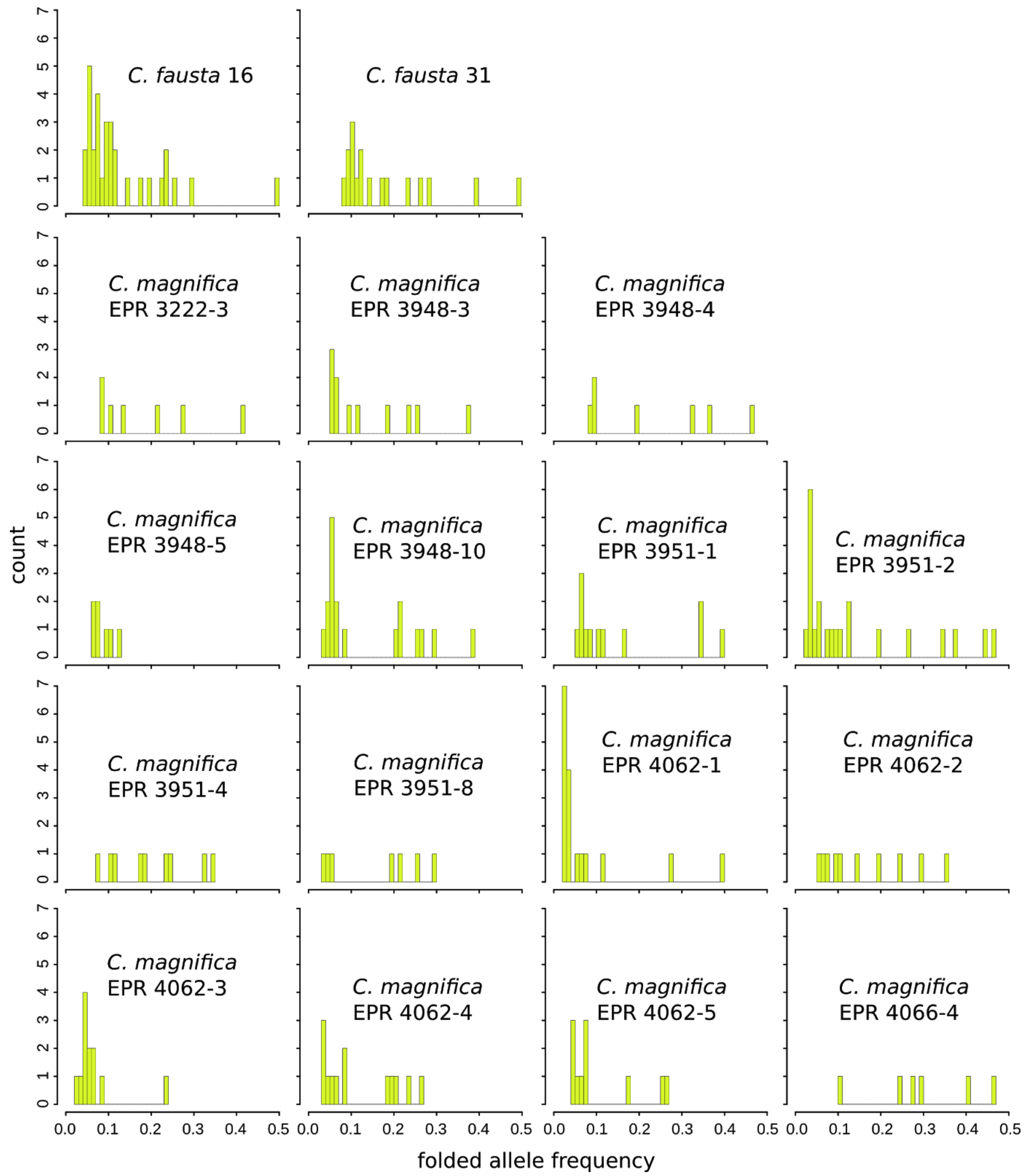


**S4 Fig.** Within-host folded allele frequency spectra for all *Calyptogena fausta* and *Calyptogena magnifica* intrahost samples with at least 50x Illumina sequencing coverage (see S1 Table for coverages and S5 Table for diversity statistics).

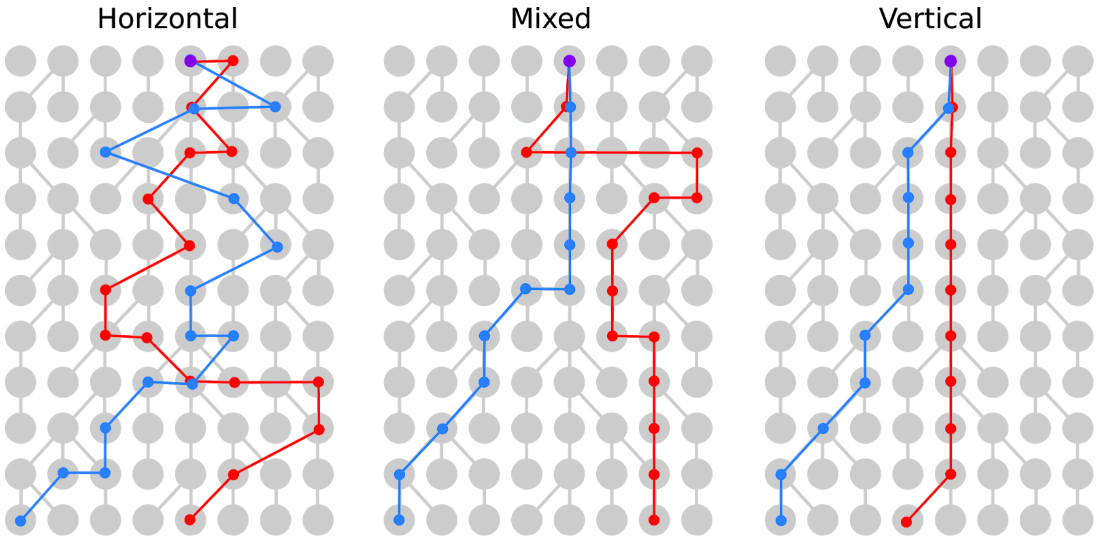


**S5 Fig.** Endosymbiont inheritance modes. Our generalized coalescent model of endosymbiont inheritance includes symbiont transmission modes ranging from strict horizontal transmission to strict vertical transmission, with mixed modes, exhibiting both horizontal and vertical strategies, in between. The host populations (grey) undergo Wright-Fisher reproduction. Endosymbiont lineages (red and blue) either switch between host lineages or are inherited, depending on the transmission mode, until they coalesce in the same host lineage (purple).

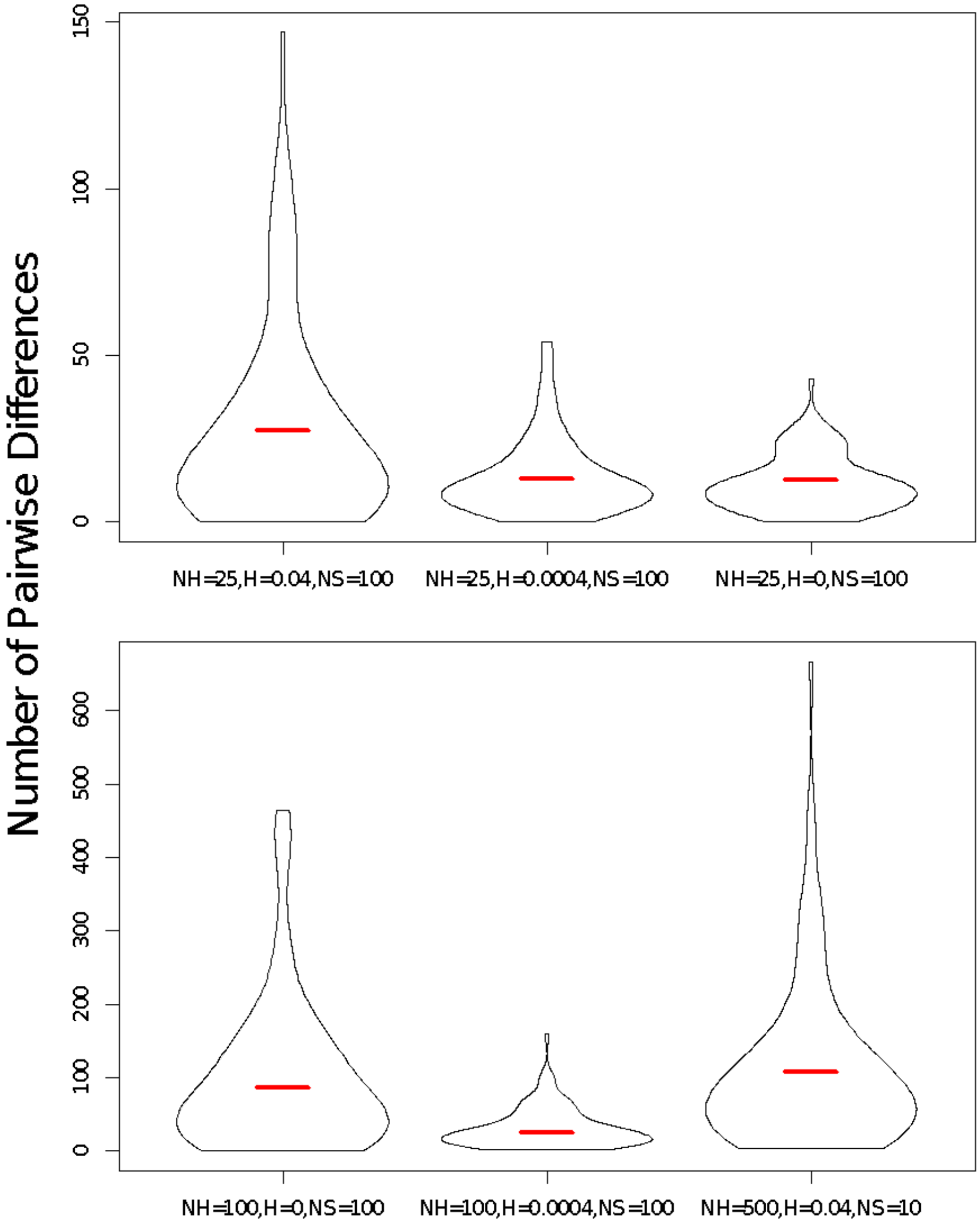


**S6 Fig.** The observed number of pairwise differences across a range of parameters under the endosymbiont population model described above. Each distribution is 100 replicates with varying $N_H$, H, and $N_S$. The expectation following Equation 9 above is plotted as a red line and differs by less than 2 segregating sites from the observed mean for all cases investigated here.

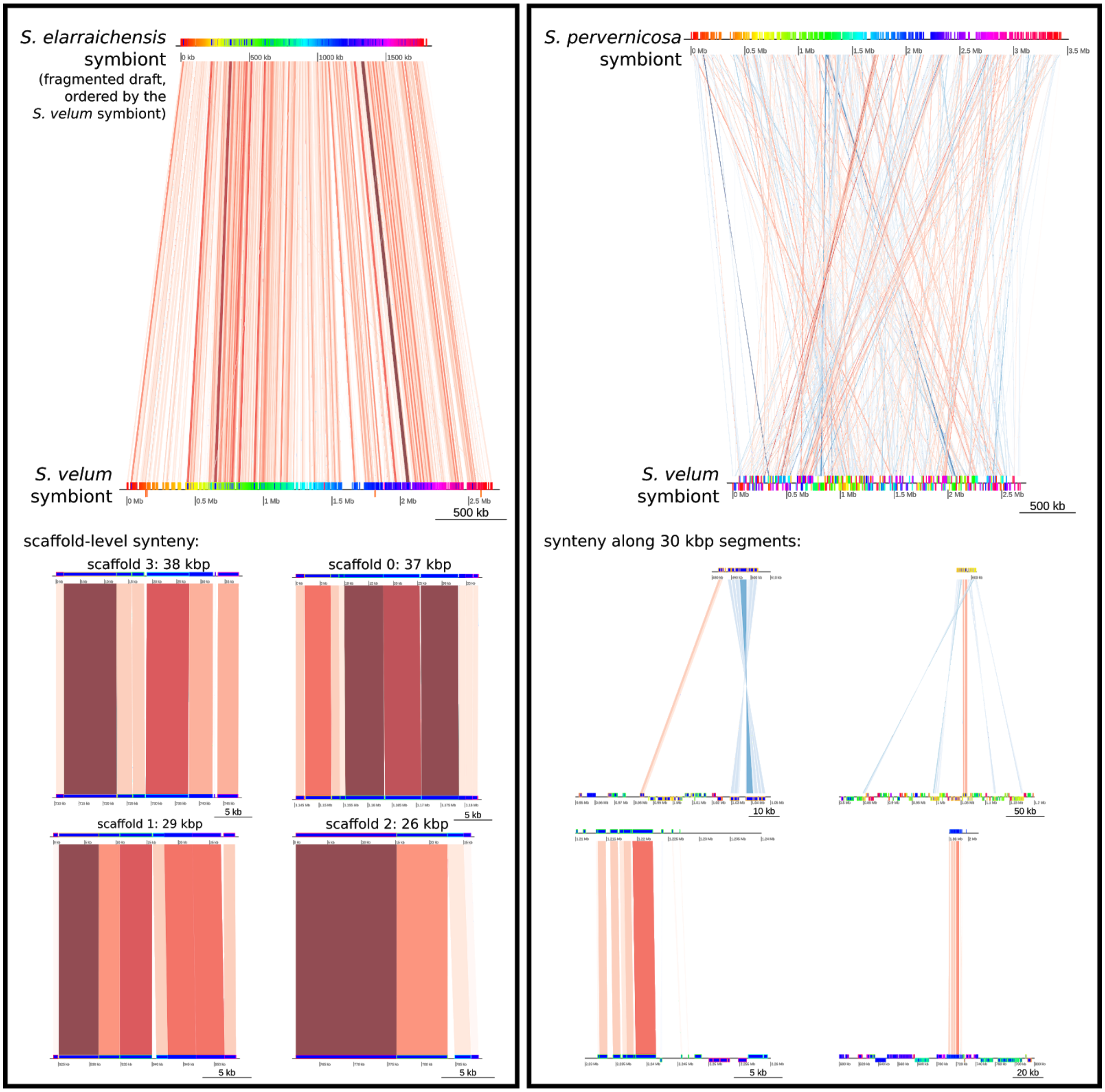


**S7 Fig.** Local alignments suggest that few rearrangements have occurred between the *S. velum* and *S. elarraichensis* symbiont genomes. *S. elarraichensis* symbiont appears to be the closest known relative of the *S. velum* symbiont, however material is exceedingly hard to obtain for this association, which occurs at a mud volcano at approximately 500-1000 m depth, and only a fragmented draft genome assembly was available. However, even these relatively short range segments reveal complete synteny (left). In comparison, over the same genomic distances, many rearrangements are evident between *S. velum* and *S. pervernicosa* (right), with the minority of segments retaining synteny.

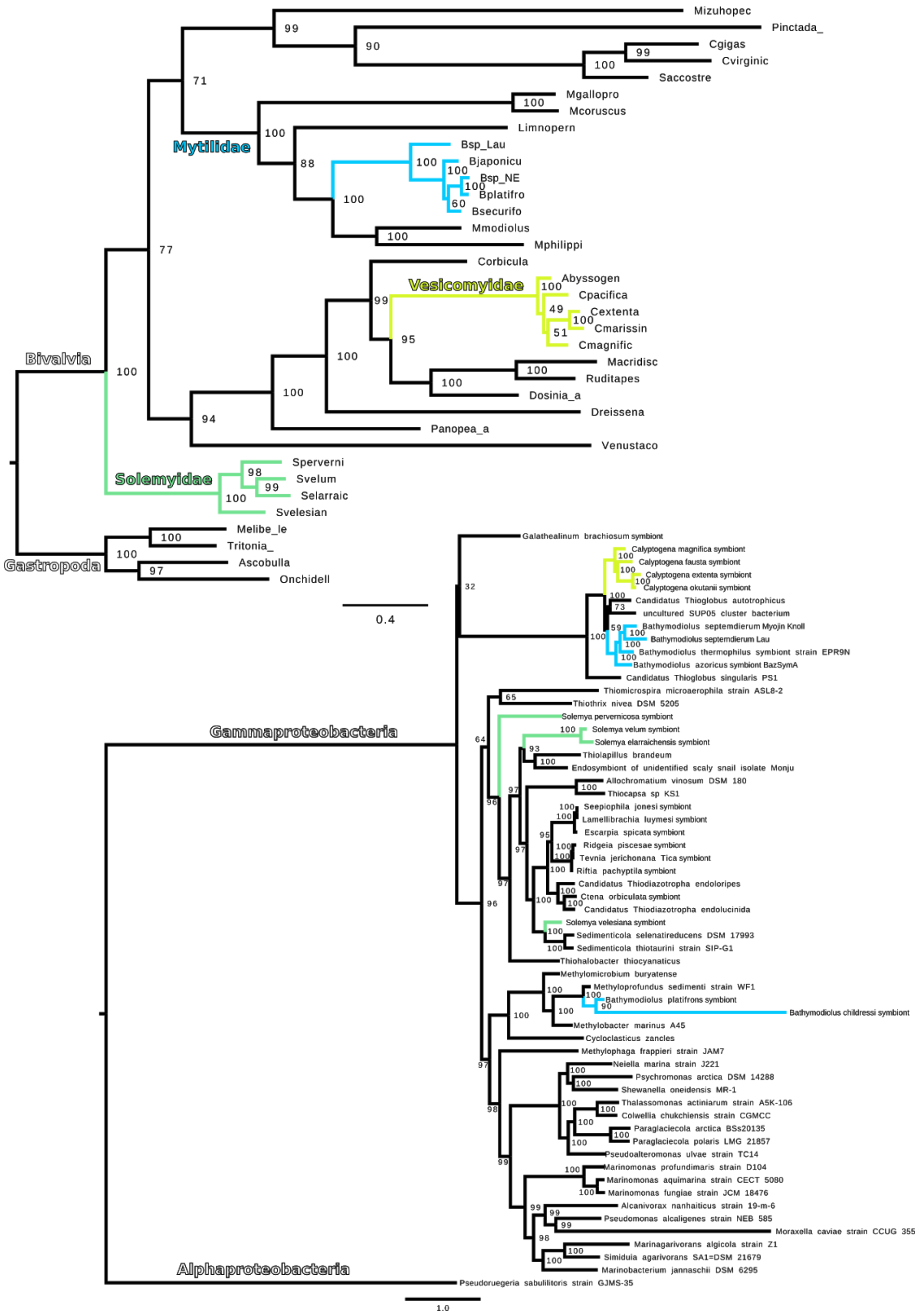


**S8 Fig.** Maximum likelihood phylogenies for host mitochondria (top) and symbionts (bottom). Groups of chemosynthetic associations are colored as in Fig. 1: yellow = vesicomyids, green = solemyids, and blue = bathymodiolids. Mitochondrial and symbiont trees are rooted by gastropod and alphaproteobacterial outgroups, respectively. Scale bar = substitutions per site. Bootstrap support values indicated at nodes.

#### Supporting Tables

**See Excel file: TableS1-SequencingStats.xlsx**

**S1 Table.** Sample, sequencing library, and mapping coverage information. The second set of coverages listed for *C. fausta* apply to the libraries used for the intra-host analysis.

| | symbiont | | | | | | | | | | | mitochondrian | | |
|---|---|---|---|---|---|---|---|---|---|---|---|---|---|---|
| species | scaffold count | total length (Mbp) | longest scaffold (Mbp) | N50 (Mbp) | % GC | gene count | % complete (34 "essential genes") | ribosome operon count | CheckM completeness (%) | CheckM contamination (%) | CheckM strain heterogeneity (%) | length (Kbp) | % GC | gene count |
| *Bathymodiolus septemdierum* | 24 | 2.37 | 1.33 | 1.33 | 36 | 3112 | 94.12 | 1 | 92.42 | 2.32 | 83.33 | 17.19 | 55 | 38 |
| *Bathymodiolus childressi* | 207 | 1.99 | 0.04 | 0.01 | 41 | 3287 | 76.47 | 1 | 43 | 0.34 | 0 | 17.96 | 55 | 43 |
| *Solemya pervernicosa* | 1 | 3.45 | 3.45 | 3.45 | 54 | 4194 | 97.06 | 1 | 97.34 | 0.69 | 0 | 16.55 | 51 | 37 |
| *Calyptogena fausta* | 1 | 1.19 | 1.19 | 1.19 | 37 | 1371 | 97.06 | 1 | 98.05 | 0 | 0 | 17.08 | 50 | 39 |
| *Calyptogena extenta* | 2 | 1.02 | 0.60 | 0.60 | 31 | 1090 | 100 | 1 | 92.73 | 0 | 0 | 17.81 | 50 | 38 |

**S2 Table.** *De novo* reference assemblies were assembled with Nanopore reads and polished with Illumina data (realigned coverage not shown). Illumina reads were used for individual sample genotype calling. There were no gaps (Ns) in any of the assemblies. The percent complete measure reflects how many of the 34 "essential genes" (see Materials and Methods) were found in the assembled genomes.

| **host family** | **host species** | **symbiont phylum** | **symbiont species** | **reference/rationale for species name** |
|---|---|---|---|---|
| Mytilidae | *Bathymodiolus septemdierum* (from Lau basin) | Gammaproteobacteria | *Candidatus Thiodubiliella endoseptemdiera sp. nov.* | this study; genus named after the chemosynthetic energy source for this symbiont (Thio=sulfide) and Prof. Nicole Dubilier who has worked on the bathymodiolin deep-sea symbioses throughout her career and contributed significant knowledge to the field about their biology; species named after host species, with the "endo" prefix representing the symbiotic association |
| Mytilidae | *Bathymodiolus childressi* (from New England) | Gammaproteobacteria | *Candidatus Methanofishera endochildressiae sp. nov.* | this study; genus named after the chemosynthetic energy source for this symbiont (Methano=methane) and Prof. Charles Fisher who has worked on the bathymodiolin deep-sea symbioses throughout his career and contributed significant knowledge to the field about their biology; species named after host species, with the "endo" prefix representing the symbiotic association |
| Solemyidae | *Solemya velum* | Gammaproteobacteria | *Candidatus Cavanaughia endovela sp. nov.* | this study; genus named after Prof. Colleen Cavanaugh who was one of the first to discover chemosynthetic symbiosis and worked on S. velum throughout her career and contributed significant knowledge to the field about their biology; species named after host species, with the "endo" prefix representing the symbiotic association |
| Solemyidae | *Solemya pervernicosa* (formerly *reidi*) | Gammaproteobacteria | *Candidatus Reidiella endopervernicosa sp. nov.* | this study; genus named after Prof. Robert Reid who first reported on these mouthless and gutless protobranch bivalves (from the synonymized *S. reidi*); species named after host species, with the "endo" prefix representing the symbiotic association |
| Vesicomyidae | *Calyptogena fausta* | Gammaproteobacteria | *Candidatus Ruthia endofausta sp. nov.* | this study; placed in genus named for other "group 2" vesicomyid symbiont (Kuwahara et al. 2011); species named after host species, with the "endo" prefix representing the symbiotic association |
| Vesicomyidae | *Calyptogena* (formerly *Ectenagena*) *extenta* | Gammaproteobacteria | *Candidatus Vesicomyosocius endoextente sp. nov.* | this study; placed in genus named for other "group 1" vesicomyid symbiont (Kuwahara et al. 2011); species named after host species, with the "endo" prefix representing the symbiotic association |
| Vesicomyidae | *Phreagena* (formerly *Calyptogena*) *okutanii* | Gammaproteobacteria | *Candidatus Vesicomyosocius okutanii* | Kuwahara et al. 2007 *Current Biology* |
| Vesicomyidae | *Calyptogena magnifica* | Gammaproteobacteria | *Candidatus Ruthia magnifica* | Newton et al. 2007 *Science* |

**S3 Table**. Symbiont species named in this study and named previously. See S3 Supporting Text for full diagnoses and descriptions.

| host species | symbiont | | | | | | | | | | | mitochondrian | | | |
|---|---|---|---|---|---|---|---|---|---|---|---|---|---|---|---|
| | segregating sites (S) | pairwise differences (pi) | Watterson's Theta | Tajima's D | RF estimate theta (Nμ) | 95% CI theta (Nμ) | Fst | 95% CI Fst | RF estimate (rho*l) | 95% CI (rho*l) | rho*l/ theta | segregating sites (S) | pairwise differences (pi) | Watterson's Theta | Tajima's D |
| *Bathymodiolus septemdierum* | 12,565 | 3.32E-03 | 2.89E-03 | 0.64 | 3.17E-03 | 3.01E-03 - 3.33E-03 | 6.17E-02 | 0.057 - 0.067 | 22.52 | 3.09 - 164.03 | 7114 | 327 | 2.88E-03 | 5.21E-03 | -1.90 |
| *Bathymodiolus childressi* | 28,199 | 2.89E-03 | 3.09E-03 | -0.31 | 2.91E-03 | 2.79E-03 - 3.04E-03 | 9.49E-02 | 0.08 - 0.114 | 46.29 | 27.82 - 77.02 | 15896 | 240 | 4.19E-03 | 5.58E-03 | -1.19 |
| *Solemya velum* | 1,722 | 1.66E-04 | 1.58E-04 | 0.20 | 1.61E-04 | 1.56E-04 - 1.65E-04 | 5.55E-02 | 0.04 - 0.072 | 17.34 | 2.57 - 117.16 | 107881 | 75 | 9.19E-04 | 1.28E-03 | -1.15 |
| *Solemya pervernicosa* | 93,013 | 1.44E-02 | 1.25E-02 | 1.01 | 8.37E-03 | 7.78E-03 - 8.99E-03 | 1.36E-01 | 0.052 - 0.266 | 8.69 | 3.13 - 24.14 | 1038 | 73 | 2.93E-03 | 3.03E-03 | -0.20 |
| *Calyptogena magnifica* | 781 | 9.86E-05 | 1.82E-04 | -1.98 | 9.11E-05 | 8.58E-05 - 9.68E-05 | 9.99E-01 | 0.998 - 1 | 0.42 | 0.15 - 2.4 | 4629 | 31 | 3.47E-04 | 4.79E-04 | -1.11 |
| *Calyptogena fausta* | 815 | 1.19E-04 | 2.00E-04 | -2.03 | 1.30E-04 | 1.34E-04 - 1.38E-04 | 9.00E-01 | 0.812 - 0.963 | 0.47 | 0.11 - 1.95 | 3625 | 73 | 1.42E-03 | 1.43E-03 | -0.03 |

**S4 Table.** Between-host symbiont population statistics calculated from consensus symbiont and mitochondrial genome sequences. Random Forest (RF) theta and log10(rho*l) estimates were inferred by fitting genome-wide values of pi, Watterson's Theta, and 4-gamete sites to values generated in coalescent simulations.

| | | | symbionts | | mitochondria | |
|---|---|---|---|---|---|---|
| Host family | Host species | Sample | within host S (SNPs, INDELS) | within host average pi | heteroplasmy S (SNPs, INDELS) | heteroplasmy average pi |
| Mytilidae | ***Bathymodiolus septemdierum* from Lau Basin** | BathyLao_211gill | 47334 (45588, 1746) | 5.25E-03 | 8 (4, 4) | 4.44E-05 |
| | | BathyLao_217gill | 59243 (57402, 1841) | 5.95E-03 | 2 (1, 1) | 3.25E-06 |
| | | BathyLao_219gill | 56323 (54439, 1884) | 5.54E-03 | 3 (1, 2) | 4.76E-06 |
| | | BathyLao_221gill | 64204 (62231, 1973) | 6.27E-03 | 2 (1, 1) | 6.14E-06 |
| | | BathyLao_227gill | 59234 (57353, 1881) | 5.90E-03 | 2, (1, 1) | 7.17E-06 |
| | | BathyLao_229gill | 55406 (53628, 1778) | 5.39E-03 | 2 (1, 1) | 4.68E-06 |
| | | BathyLao_231gill | 65133 (63058, 2075) | 6.29E-03 | 0 | 0.00E+00 |
| | | BathyLao_233gill | 55330 (53584, 1746) | 5.27E-03 | 3 (1, 2) | 4.49E-06 |
| | | BathyLao_239gill | 54328 (52621, 1707) | 5.38E-03 | 2 (1, 1) | 6.28E-06 |
| | | BathyLao_243gill | 46996 (45350, 1646) | 4.81E-03 | 4 (2, 2) | 1.14E-05 |
| | | BathyLao_443gill | 60208 (58405, 1803) | 6.09E-03 | 0 | 0.00E+00 |
| | | BathyLao_445gill | 56210 (54432, 1778) | 5.24E-03 | 4 (2, 2) | 1.86E-05 |
| | | BathyLao_447gill | 57827 (56025, 1802) | 5.56E-03 | 4 (2, 2) | 1.06E-05 |
| | | BathyLao_451gill | 65165 (63213, 1952) | 6.34E-03 | 1 (0,1) | 0.00E+00 |
| | | BathyLao_455gill | 39323 (37749, 1574) | 4.73E-03 | 1 (0,1) | 0.00E+00 |
| | | BathyLao_457gill | 38104 (36595, 1509) | 4.40E-03 | 1 (0,1) | 0.00E+00 |
| | | BathyLao_459gill | 44755 (43094, 1661) | 5.27E-03 | 1 (0,1) | 0.00E+00 |
| | ***Bathymodiolus childressi* from New England** | BathyNE_4727-3 | 1173 (694, 479) | 1.64E-04 | too low depth | too low depth |
| | | BathyNE_4727-4 | 1152 (668, 484) | 1.58E-04 | too low depth | too low depth |
| | | BathyNE_4727-5 | 2460 (1598, 862) | 3.58E-04 | 2 (0, 2) | 0.00E+00 |
| | | BathyNE_4727-9 | 240 (78, 162) | 2.02E-05 | too low depth | too low depth |
| | | BathyNE_4727-10 | 2211 (1479, 732) | 3.27E-04 | too low depth | too low depth |

| | | | | | | |
|---|---|---|---|---|---|---|
| | | BathyNE_4727-11 | 16693 (10694, 5999) | 1.93E-03 | 6 (4, 2) | 5.87E-05 |
| | | BathyNE_4728-2 | 6025 (4038, 1987) | 8.10E-04 | 1 (0, 1) | 0.00E+00 |
| | | BathyNE_4728-4 | 14766 (12109, 2657) | 2.19E-03 | 2 (1, 1) | 1.25E-05 |
| | | BathyNE_4728-9 | 2105 (1419, 686) | 3.10E-04 | 0 (0, 0) | 0.00E+00 |
| | | BathyNE_4728-12 | 5533 (4197, 1336) | 8.42E-04 | 1 (0, 1) | 0.00E+00 |
| | | BathyNE_4728-13 | 6454 (3840, 2614) | 7.95E-04 | 0 (0, 0) | 0.00E+00 |
| **Solemyidae** | ***Solemya velum* from Rhode Island** | RI36gill | 670 (558, 112) | 2.92E-05 | 2 (1, 1) | 8.51E-06 |
| | | RI37gill | 478 (372, 106) | 2.77E-05 | 2 (1, 1) | 6.69E-06 |
| | | RI38gill | 829 (728, 101) | 5.78E-05 | 0 | 0.00E+00 |
| | | RI39gill | 3414 (3273, 141) | 2.07E-04 | 4 (2, 2) | 1.84E-05 |
| | | RI41gill | 83 (7, 76) | 1.41E-06 | 0 | 0.00E+00 |
| | | RI44gill | 362 (229, 133) | 2.37E-05 | 2 (1, 1) | 5.37E-06 |
| | | RI47gill | 463 (330, 133) | 3.25E-05 | 0 (0, 1) | 0.00E+00 |
| | | RI48gill | 283 (144, 139) | 2.35E-05 | 3 (2, 1) | 4.12E-05 |
| | | RI50gill | 234 (112, 122) | 1.67E-05 | 1 (0, 1) | 0.00E+00 |
| | | RI51gill | 513 (382, 131) | 4.29E-05 | 1 (0, 1) | 0.00E+00 |
| | | RI53gill | 762 (642, 120) | 7.02E-05 | 2 (1, 1) | 5.75E-06 |
| | | RI55gill | 607 (439, 168) | 7.83E-05 | 3 (1, 2) | 1.34E-05 |
| | | RI56gill | 531 (383, 148) | 6.81E-05 | 1 (0, 1) | 0.00E+00 |
| | | RI57gill | 197 (68, 129) | 1.27E-05 | 3 (0, 3) | 0.00E+00 |
| | | RI58gill | 168 (46, 122) | 8.72E-06 | 4 (3, 1) | 8.74E-05 |
| | | RI59gill | 281 (168, 113) | 3.10E-05 | 1 (0, 1) | 0.00E+00 |
| | | RI60gill | 130 (19, 111) | 3.68E-06 | 1 (0, 1) | 0.00E+00 |
| | | RI61gill | 401 (267, 134) | 3.14E-05 | 5 (2, 3) | 2.63E-05 |
| | | RI62gill | 449 (341, 108) | 4.48E-05 | 1 (0, 1) | 0.00E+00 |
| | | RI63gill | 533 (387, 146) | 7.07E-05 | 2 (1, 1) | 1.73E-05 |
| | ***Solemya pervernicosa* from California** | Spervernicosa_2gill | 73925 (72871, 1054) | 1.03E-02 | 2 (5, 1) | 4.99E-06 |
| | | Spervernicosa_4gill | 78934 (77545, 1389) | 9.62E-03 | 6 (5, 1) | 3.16E-05 |
| | | Spervernicosa_5gill | 20658 (19987, 671) | 3.02E-03 | 6 (3, 3) | 7.33E-05 |
| | | Spervernicosa_6gill | 308 (191, 117) | 1.49E-05 | 6 (6, 0) | 3.70E-05 |
| | | Spervernicosa_7gill | 76871 (75499, 1372) | 1.08E-02 | 6 (5, 1) | 8.48E-05 |
| | | Spervernicosa_8gill | 85328 (84237, 1091) | 9.29E-03 | 8 (6, 2) | 7.46E-05 |

| | | | | | | |
|---|---|---|---|---|---|---|
| Vesicomyidae | ***Calyptogena magnifica* from the East Pacific Rise** | CmagEPR3222_3 gill | 18 (7, 11) | 1.70E-06 | 7 (6, 1) | 1.48E-04 |
| | | CmagEPR3948_3 gill | 22 (11, 11) | 2.13E-06 | 7 (5, 2) | 1.30E-04 |
| | | CmagEPR3948_4 gill | 17 (7, 10) | 1.95E-06 | 8 (7, 1) | 1.82E-04 |
| | | CmagEPR3948_5 gill | 14 (7, 7) | 9.58E-07 | 4 (1, 3) | 2.67E-05 |
| | | CmagEPR3948_1 0gill | 35 (18, 17) | 3.32E-06 | 12 (11, 1) | 2.38E-04 |
| | | CmagEPR3951_1 gill | 26 (12, 14) | 2.48E-06 | 1 (1, 0) | 2.73E-05 |
| | | CmagEPR3951_2 gill | 42 (22, 20) | 3.91E-06 | 10 (10, 0) | 2.55E-04 |
| | | CmagEPR3951_3 gill | 3 (0, 3) | 0.00E+00 | 0 | 0.00E+00 |
| | | CmagEPR3951_4 gill | 19 (9, 10) | 2.39E-06 | 0 | 0.00E+00 |
| | | CmagEPR3951_6 gill | 14 (4, 10) | 1.57E-06 | 0 | 0.00E+00 |
| | | CmagEPR3951_8 gill | 15 (7, 8) | 1.47E-06 | 2 (0, 2) | 0.00E+00 |
| | | CmagEPR4062_1 gill | 33 (17, 16) | 1.79E-06 | 27 (25, 2) | 4.77E-04 |
| | | CmagEPR4062_2 gill | 21 (10, 11) | 2.20E-06 | 3 (1, 2) | 2.61E-05 |
| | | CmagEPR4062_3 gill | 23 (12, 11) | 1.23E-06 | 22 (18, 4) | 3.43E-04 |
| | | CmagEPR4062_4 gill | 25 (13, 12) | 2.16E-06 | 0 | 0.00E+00 |
| | | CmagEPR4062_5 gill | 22 (11, 11) | 1.67E-06 | 3 (1, 2) | 2.38E-05 |
| | | CmagEPR4066_4 gill | 18 (6, 12) | 2.08E-06 | 1 (0, 1) | 0.00E+00 |
| | ***Calyptogena fausta* from the Juan de Fuca Ridge** | Cfausta_16gill | 85 (31, 54) | 5.42E-06 | 1 (0, 1) | 0.00E+00 |
| | | Cfausta_18gill | 138 (84, 54) | 2.40E-05 | 1 (0, 1) | 0.00E+00 |
| | | Cfausta_31gill | 73 (17, 56) | 3.99E-06 | 0 (0, 0) | 0.00E+00 |
| | | Cfausta_32gill | 2 (0, 2) | 0.00E+00 | too low depth | too low depth |

**S5 Table.** Within-host symbiont and mitochondrial genetic diversity statistics. Mapping coverages in S1 Table.

| sample size | Oob score rho*l | Oob score theta |
|---|---|---|
| 6 | 0.88 | 0.93 |
| 10 | 0.91 | 0.95 |
| 17 | 0.93 | 0.97 |
| 18 | 0.95 | 0.98 |
| 20 | 0.96 | 0.98 |

**S6 Table.** Out-of-bag (oob) scores for random-forest models for each parameter of interest, rho*l and theta, and for each sample size of endosymbiont individuals considered. Oob scores indicate how often the trained model is able to predict known values, with perfect prediction equal to one.

| | **Proportion 4-gametes found from WIthin-Host** | | | **Proportion 4-gametes found from consensus** | | |
|---|---|---|---|---|---|---|
| **Host Species** | **10 bp** | **100 bp** | **1000 bp** | **10 bp** | **100 bp** | **1000 bp** |
| *S. velum* | 0.350 | 0.559 | 0.678 | 0.261 | 0.474 | 0.588 |
| *S. pervernicosa* | 0.096 | 0.122 | 0.231 | 0.000 | 0.105 | 0.206 |
| *B. septemdierum str. Lau* | 0.136 | 0.207 | 0.560 | 0.015 | 0.200 | 0.401 |
| *B. childressi str. NE* | 0.583 | 0.753 | 0.795 | 0.465 | 0.651 | 0.656 |

**S7 Table.** Proportion of within-host variant sites that pass the 4-gamete test for recombination based upon read and read pair data over the given genomic intervals (constrained by Illumina library insert sizes).

| **host species** | **genome size (bp)** | **gene count** | **bp / gene** | **average GC content (%)** | **# ME regions >=90% identity** |
|---|---|---|---|---|---|
| *B. septemdierum* | 2370579 | 3112 | 761.75 | 36.28 | 4 |
| *B. childressi* | 1990083 | 3287 | 618.34 | 40.70 | 6 |
| *S. velum* | 2672015 | 2741 | 974.83 | 50.46 | 9 |

| | | | | | |
|---|---|---|---|---|---|
| *S. pervernicosa* | 3446950 | 4194 | 821.88 | 54.10 | 10 |
| *C. magnifica* | 1160782 | 2176 | 533.45 | 34.03 | 5 |
| *C. fausta* | 1188809 | 1371 | 867.11 | 36.69 | 4 |
| *C. extenta* | 1021947 | 1090 | 937.57 | 31.13 | 2 |

**S8 Table.** Comparative symbiont genome statistics and mobile element (ME) content. MEs were identified as regions within the symbiont genomes with high sequence identity to elements in insertion sequence, phage, and integrative conjugative element databases.

**See Excel file: S9Table-SymbiontMobileElementHits**
**S9 Table.** Full list of ICEberg and ACLAME database mobile element hits with >=90% sequence identity to endosymbiont genomic regions and genes.

**See Excel file: TableS10-taxa_for_mitochondrial_dating**
**S10 Table.** Taxa and accession numbers used in the mitochondrial genome phylogenetic analysis and divergence dating.

**See Excel file: TableS11-taxa_for_bacterial_dating**
**S11 Table.** Taxa and accession numbers used in the bacterial whole genome phylogenetic analysis and divergence dating.

**See Excel file: TableS12-divergence_date_estimates**
**S12 Table.** Divergence date estimates from different Beast2, TimeTree, and PATHd8 runs with different parameter values.
